# Small extracellular vesicles from iPSC-MSC lose their regenerative potential upon UV-C irradiation

**DOI:** 10.1101/2021.09.27.461979

**Authors:** M.C. Biani, A. Lombardi, A. Norris, P. Bucci, A. La Greca, A. Waisman, L.N. Moro, G. Sevlever, J. Montanari, S. Miriuka, C. Luzzani

## Abstract

Mesenchymal Stem Cells derived from induced Pluripotent Stem cells (iPSC-MSC) have become a promising alternative to classical Mesenchymal Stem Cells in regenerative medicine. Their properties -as immunomodulatory and regenerative capacities-are in part due to the secretion of Extracellular Vesicles (EVs). Small EVs (sEVs) with sizes that range from 50 to 120 nm contain proteins, lipids, and nucleic acids that exert a role in cellular communication. Their content will depend on the cell of origin and its physiological state, thus the message they convey might change in response to changes in cellular conditions. In particular, the DNA damage response (DDR) has been reported to modulate sEVs secretion. In this work, we analyze how UV-C radiation upon iPSC-MSC alter sEVs secretion, cargo and bystander effect. Here, we confirm that UV-C radiation causes DDR in a dose dependent manner. In addition, we found that UV-C induced stress did not modulate the expression of genes that participate in sEVs biogenesis pathway. Consequently, we found that the amount of sEVs secreted by radiated and non-irradiated cells remained stable. However, sEVs from radiated cells were unable to promote cell migration in their target cells. Moreover, a label-free proteomic analysis revealed that UV-C induced DDR produces sEVs with an altered cargo, rich in migration-inhibiting proteins, and resulting in a less stromal-oriented repertoire.

## Introduction

Mesenchymal Stem/Stromal Cells (MSC) are currently being tested for several clinical applications (www.clinicaltrials.gov). They are generally considered as safe and their immuno-modulatory and regenerative capacities make them a perfect alternative for pathologies that involve alterations in the body’s immune response (1, 2). However, MSC massive use at the bedside continues to pose some doubts since several hurdles such as limited life span and rapid decay in their therapeutic properties in culture remain unsolved (3, 4). A potential solution to those drawbacks is the isolation of MSC from Pluripotent Stem Cells (PSC) (5, 6). Unlike MSC, PSC are able to differentiate to cells of the three lineages, endoderm, mesoderm and ectoderm. Induced Pluripotent Stem Cell-Derived MSC (iPSC-MSC) are cells that resemble MSC in all its characteristics, namely molecular markers, differentiation capacity and immuno-modulatory properties, and can be readily obtained in a clinical compliant and inexpensive manner over a 30-day differentiation protocol (7).

Both immuno-modulatory and regenerative properties of MSC are partially explained by the secretion of soluble factors and extracellular vesicles (EVs) (8–11). Extracellular vesicles are particles released by most types of cells that are delimited by a lipid bilayer and may enclose DNA, miRNA, RNA and proteins. They were usually classified according to their size and cargo roughly into exosomes, microvesicles, and apoptosomes. While microvesicles are large EVs that bud directly from the plasma membrane, exosomes are small EVs (sEVs) with sizes that range from 50 to 120 nm and have an endosomal origin. This classification, however, is presently challenged by evidence that points to heterogeneity in EVs composition and origin within the same range of size distribution (12). Hence, in accordance with MISEV2018 (13), we will hereon use the term sEVs to name particles up to 120 nm that are positive for the exosomal markers CD63, CD9, and CD81.

Though currently known to be key mediators in cell-cell communication, EVs were once thought to be a means of disposing of cell’s toxic material (14, 15). However, the message they convey might change in response to altered cellular conditions, an effect known as the bystander effect. In particular, DNA damage seems to have profound consequences on EVs biogenesis and their effect on target cells.

Cells are particularly sensitive to UV-C radiation (190-280 nm) since their wavelength is within the absorption range of DNA, RNA, and proteins (16). Within DNA lesions, double-strand breaks (DSBs) can compromise histone integrity and consequently its function. The histone H2 variant, histone 2AX (H2AX) suffers S-139 phosphorylation caused by the DNA damage response (DDR) (16) in which ATM and ATR kinases are the main damage sensors (17). The purpose of this mechanism is the efficient activation of the intra-S check-point response to arrest the cell cycle and activate extensive cellular networks to repair the damage. Another central node of the damage response is p53, whose activation leads to cell cycle arrest allowing time for damage repair or induction of apoptosis to prevent the division of damaged cells (18).

EVs biogenesis seems to be highly influenced by DNA damage. On the one hand, senescent cells were reported to increase EVs secretion, professedly via a p53-dependent secretory pathway. In this model, activation of p53 is proposed to lead to transcriptional induction of STEAP3 gene (also known as TSAP6) which, in turn, would augment a non-canonical secretory pathway (19–21). Though an increase in exosome secretion was confirmed in immortalized keratinocyte HaCaT (22), breast carcinoma MCF7 (23), and human prostate cancer cell lines (24), the mechanism by which Steap3 regulates EV secretion is still poorly understood. On the other hand, EVs secreted from ionizing-radiated cells suffer changes in their composition and induce DNA and chromosomal damage and promote cell death in recipient cells (19). In addition, EVs secreted by senescent cells were shown to have a different proteomic profile compared to non-senescent cells, stimulating the proliferation of several types of cancer cells (25). Last, DNA damage also seems to modulate toxic DNA loading into sEVs (15).

Extracellular vesicles have the potential to generate molecular changes in their target cells allowing a coordinated response between adjacent cells. Disrupting the cellular homeostasis by generating genomic stress by UV-C radiation provides a means to study how this transmitted message can change. In this work we focus on how DNA damage by UV-C radiation may modulate sEVs secretion and cargo, altering MSC EVs usual pro-regenerative effect over the cellular stroma. Here we report that this type of radiation does not seem to modulate the expression of EVs biogenesis-related genes nor significantly change the amount of EVs secreted by MSC. However, UV-C irradiation does diminish sEVs ability to promote migration over target cells. Moreover, we found that this loss of function is, at least in part, explained by a profound change in EV proteomic composition.

## Methods

### Cell culture

Mesenchymal Stem Cells (MSC) were derived from Induced Pluripotent Stem Cells (iPSC) using a differentiation protocol previously published by our laboratory (7). Briefly, on day 0 of differentiation PSCs were incubated with Accutase until the cells were completely dissociated, and plated onto Geltrex-coated plates. Alpha-minimum essential medium (α-mem) supplemented with platelet lysate 10% and B27 1/100 (all from Life Technologies, Carlsbad, CA, USA). A ROCK inhibitor (Y27632 10 ng/ml; Tocris, Avon, Bristol, UK) was added every time the cells were passaged until day 14 of differentiation. From that day onwards and during 14 more days, the cells were frequently passed on plastic dishes with no coating and were grown in medium supplemented with PL 10% and penicillin-streptomycin until a complete mesenchymal phenotype was reached. Wharton-Jelly Mesenchymal Stem Cells (WJ-MSC) were grown in α-mem supplemented with PL 10%. All experiments were performed using MSCs maintained in culture until passage ten or less.

### iPSC-MSC UV-C radiation and media conditioning

iPSC-MSC were seeded to reach 80% of confluence using α-mem supplemented with PL 10%. The cells were then washed with PBS and irradiated with three intensities of UV-C (0.001 J/cm2, 0.01 J/cm2 and 0.1 J/cm2) using a CL-1000 Ultraviolet crosslinker (254 nm, UVP, Upland, CA, USA) with the culture plate uncovered and without culture medium in order to maximize the radiation efficiency. E8-defined medium (Life Technologies, Carlsbad, CA, USA) or α-mem supplemented with 10% PL (depending on the experiment) was added to the culture plates after irradiation.

### EV isolation

EVs were isolated by Size Exclusion Chromatography using a qEV2 Size Exclusion Column (iZON science, New Zealand) according to manufacturer instructions. Briefly, cells were seeded to reach 80% of confluence and cultured for 24 hs in E8-defined, serum-free medium (Life Technologies, Carlsbad, CA, USA). The cell culture supernatants were then centrifuged at 4°C, 2000 x g for 15 min, which was followed by another centrifugation at 4°C, 10.000 x g for 30 min. Finaly, a last concentration step was performed using a 100 kDA filter.(Amicon Ultra-15, Millipore, Merck, Ireland). The centrifuged and concentrated samples were loaded into the loading frit of the reservoir and once they entered the column, the reservoir was topped up with filtered PBS. Fractions of 1.5 ml were collected and the presence of sEVs in the fractions was evidenced by Flow Cytometry.

### Quantitative qPCR analysis

Total RNA from cell extracts was isolated using Trizol (Life Technologies, Carlsbad, CA, USA) as described by the manufacturer. Total RNA was retro-transcribed using MMLV (Promega, Madison, Wisconsin, USA) and random primers (Life Technologies, Frederick, MD, USA). Quantitative RT-PCR was performed using the FastStart Universal SYBR Green Master (ROX) (Roche, Mannheim, Germany) in a Step One Plus Real-Time PCR system (Applied Biosystems, Foster City, CA, USA). EVs containing fractions from 24 h conditioned medium were concentrated using a 100 kDA filter (Amicon Ultra-15, Millipore, Merck, Irealnd) and total RNA was isolated as previously described. The list of the primers used is presented in Supplementary Table 1.

### Flow cytometry analysis

Flow cytometry analyses were performed in a BD Accuri cytometer. In order to confirm on which fractions exosomes were present, 100 ul of each fraction was bound with 5 ul of anti-CD63 antibody-coated magnetic beads (Life Technologies, Oslo, Norway), as recommended by the manufacturer. Next, anti-CD63 bound EVs were stained with an anti-CD81-PE-conjugated antibody and an anti-CD9-APC-conjugated antibody (Molecular Probes, Life Technologies, Eugene, OR, USA) for 30 min at room temperature. Beads were then washed with PBS 0.1% albumin and were analyzed. At least 10000 events per treatment were counted.

### Direct sEV counting by Tunable Resistive Pulse Sensing (TRPS)

A 1:100 dilution of the suspension of exosomes was performed prior to TRPS analysis. Then an aliquot of 40 ml was loaded into the NP200 nanopore membrane of a qViro equipment (Izon Science Ltd, Christchurch, New Zealand) with a membrane stretch between 45,66 mm and 48,66 mm. Voltage used was 0,46 V and the Average Current was 118,56 nA. Sample was calibrated with a standard of 210 nm with a concentration of 1.0 x 1020 particles per ml. Measures were taken by triplicate. A minimum of 300 particles were counted in each sample measurement.

### Wound Healing Assay

WJ-MSC were seeded to confluency in a 12-well with α-mem supplemented with 10% PL. After 14 hrs, injuries were performed manually using a micropipette tip. Immediately after this, cells were washed with PBS and the injuries were photographed to register the initial setting (time zero: 0h), followed by the addition of α-mem medium containing or not EVs collected from irradiated or non-irradiated iPSC-MSC. After 14 h, injuries were photographed to mark the conclusion of the experiment. Images were acquired via Nikon (Eclipse TS100) and were analyzed using ImageJ’s MRI Wound Healing Tool (ImageJ’s macros, open source software - Redmine repository).

### Cell proliferation assay

A cell proliferation assay was performed using CyQUANT NF Cell Proliferation Assay Kit (Invitrogen). WJ-MSC were counted and seeded at density 8000 cells per well in a 96-well plate with α-mem. After 14 h of incubation with irradiated or non-irradiated iPSC-MSC derived EVs, cells were stained following manufacturer’s indications. Briefly, growth medium was aspirated, replaced for dye solution and fluorescence at 538 nm was measured in a microplate fluorometer (Fluoroskan Ascent FL, Thermo Fisher Scientific) 30 minutes post incubation. A standard curve of fluorescence intensity versus cell number was performed in order to determine the number of cells.

### Cell cycle analysis

iPSC-MSC were seed to 80% confluency in a 12-well with α-mem and incubated during 14 h with irradiated or non-irradiated iPSC-MSC derived EVs. Then, the cells were harvested, fixed with 70% Ethanol and incubated for 2 hrs at −20°C. After this, cells were washed with PBS and resuspended in a Propidium Iodide solution (PBS, 25 μg/ml propidium iodide, 60 μg/ml Invitrogen PureLink RNAse A). After 15 minutes of dark incubation at room temperature flow cytometry analyses were performed in a BD Accuri cytometer. At least 10000 events per treatment were counted.

### Immunofluorescence microscopy

Cells were grown in 24 multiwell plates (90,000 cells per well) on coverslips previously treated 30-40 minutes with a sterile bovine gelatin solution (0.1% v/v in PBS, Sigma) and incubated for 24 hours prior to irradiation. Coverslips were transferred to a 60 mm plate, washed carefully with PBS and irradiated with UV-C light. In order to analyze the activation kinetics of different proteins, the irradiated cells were incubated for different times expressed in hours: 0 hrs (irradiated and immediately fixed), 1 hr, 3 hrs, 6 hrs, 10 hrs, 16 hrs, 24 hrs and a not irradiated control. After the different post-irradiation incubations, the cells attached to the coverslips were washed with PBS and fixed with a 4% m/v paraformaldehyde solution (PFA, Sigma) for 45 minutes at room temperature. Post fixation, three washes with a wash solution (0.1% v/v bovine serum albumin (BSA, Gibco) in PBS) were performed and then proceeded to permeate and block overnight using 0.1% v/v Triton X-100 (Sigma), 0.1% v/v BSA and 10% v/v Normal Goat Serum (NGS, Gibco) in PBS solution. After removing the permeabilization and blocking solution the coverslips were incubated in a humid chamber, overnight at 4 °C, with primary antibodies diluted in a solution of 0.1% v/v BSA, 10% v/v NGS in PBS (α-p53 (1:200, mouse monoclonal, #ab1101, Abcam); α-P-p53 Ser15 (1:200, rabbit monoclonal, #92845, Cell Signalling); α-γH2AX (1:1000, mouse monoclonal, #ab2893, Abcam)). After incubation, the coverslips were washed three times with the wash solution and incubated in humid chamber 45 minutes with secondary antibodies conjugated with fluorophores. For nuclei staining, 4,6-diamidino-2-phenylindole (1:50) (DAPI, Sigma) was added to the secondary antibody dilution. Finally, the coverslips were washed three times, mounted on slides using a mounting solution and sealed for observation under fluorescence microscope. The slides were observed with an inverted fluorescence microscope (Nikon Eclipse TE2000-S) and pictures were taken with a digital camera coupled to the microscope (Nikon DXN1200F) using the EclipseNet 1.20.0 Build 61 software. All the images were processed with ImageJ software.

### Western blot

Samples of mesenchymal stem cells irradiated with UV-C light (as mentioned before) and not irradiated were lysed 8 or 24 hours after irradiation from 100 mm culture plates with 70 μL of lysis buffer (60 mM Tris–HCl 1% SDS, pH=6,8). Lysates were boiled 3 minutes, vortexed, and boiled again for 7 minutes and then centrifuged at 13.000 rpm at room temperature for 10 minutes. The supernatants were collected. Protein concentration was determined using a commercial kit (Pierce™ BCA Protein Assay Kit, Thermo Fischer) and measured at 595 nm with a spectophotometer (BioRad iMarkTM Micro Plate Reader). Each sample was prepared using 30 μg of total proteins combined with the appropriate amount of 5X loading buffer (10% β-Mercaptoethanol) and boiled for 2 minutes. Samples were loaded on 10% SDS-polyacrylamide gels, separated by electrophoresis, and then transferred to PVDF membranes. Membranes were blocked using PBS-0.1% Tween-5% skim milk, cut based on the molecular weights indicated by the prestained marker and then incubated overnight on a shaker at 4 oC with rabbit anti-human STEAP3 (1:1000) (abcam, 151566), mouse anti-human p53 (1:1000) (abcam, 1101), and goat anti-human GAPDH (1:1000, #SC-20357, Santa Cruz). The membranes were washed with PBS-0.1% Tween three times and incubated with HRP-conjugated secondary antibodies (1:5000) 1 hour at room temperature. Finally, membranes were washed and revealed using a commercial kit (Pierce ECL Western Blotting Substrate, Thermo Fischer) and photos were taken by the ImageQuant LAS 4000 mini (GE Healthcare) camera system. All images were analyzed using ImageJ software.

### Mass spectrometry analysis

Liquid chromatography tandem mass spectrometry (LC-MS/MS) was performed on two biological replicates of non-irradiated and irradiated iPSC-MSC derived EVs. The datasets were then grouped according to the experimental condition. Each group was run at least three times. Protein digestion and mass spectrometry analysis were performed at the Proteomics Core Facility CEQUIBIEM, at the University of Buenos Aires/CONICET (National Research Council) as follows: the protein samples were reduced with dithiothreitol in 50mM of ammonium bicarbonate at a final concentration of 10mM (45 min, 56 °C) and were alkylated with iodoacetamide in the same solvent at a final concentration of 20mM (40 min, room temperature (RT), in darkness). This protein solution was precipitated with 1/5 volumes of trichloroacetic acid at 20 °C for at least 2 h and was centrifuged at maximum speed for 10 min (4 °C). The pellet was washed twice with cool acetone and was dried at RT. The proteins were resuspended in ammonium bicarbonate 50 mM, pH = 8, and were digested using trypsin (V5111; Promega, Madison, WI, USA). Next, the peptides were purified and desalted via ZipTip C18 columns (Millipore, Burlington, MA, USA). The digests were analyzed via nanoLC-MS/MS in a Thermo Scientific Q-Exactive Mass Spectrometer coupled to a nanoHPLC EASY-nLC 1000 (Thermo Scientific). For the LC-MS/MS analysis, approximately 1 μg of peptides was loaded onto the column and was eluted for 120 min using a reversed-phase column (C18, 2 μm, 100 A, 50 μm x 150 mm) Easy-Spray Column PepMap RSLC (P/NES801) that was suitable for separating protein complexes with a high degree of resolution. The flow rate used for the nanocolumn was 300 nl min1, and the solvent range used was from 7% B (5 min) to 35% B (120 min). Solvent A was 0.1% formic acid in water, whereas B was 0.1% formic acid in acetonitrile. The injection volume was 2 μL. The MS equipment has a high collision dissociation cell (HCD) for fragmentation and an Orbitrap analyzer (Q-Exactive, Thermo Scientific). A voltage of 3.5 kV was used for electrospray ionization (Thermo Scientific, EASY-SPRAY). XCalibur 3.0.63 software (Thermo Scientific) was used for data acquisition and equipment configuration that allows peptide identification simultaneously with their chromatographic separation. Full-scan mass spectra were acquired in the Orbitrap analyzer. The scanned mass range was 400–1800m/z, at a resolution of 70,000 at 400 m/z, and the 12 most intense ions in each cycle were sequentially isolated, fragmented by HCD, and measured in the Orbitrap analyzer. Peptides with a charge of +1 or with unassigned charge state were excluded from fragmentation for MS2.

### Analysis of MS data

Q-Exactive raw data was processed using Proteome Discoverer software (version 2.1.1.21, Thermo Scientific) and was searched against Homo sapiens protein sequence database with trypsin specificity and a maximum of one missed cleavage per peptide. Carbamidomethylation of cysteine residues was set as a fixed modification, and oxidation of methionine was set as variable modification. Proteome Discoverer searches were performed with a precursor mass tolerance of 10 ppm and product ion tolerance of 0.05 Da. Static modifications were set to carbamidomethylation of Cys, and dynamic modifications were set to oxidation of Met and N-terminal acetylation. Protein hits were filtered for high confidence peptide matches with a maximum protein and peptide false discovery rate of 1%, which was calculated by employing a reverse database strategy. Proteome Discoverer then calculates the Peptide Spectrum Matches (PSMs) for each protein in each condition. This parameter represents the total numbers of peptides identified for a certain protein.

### Bioinformatic analysis

A label-free semi-quantitative analysis of protein abundance was performed using the PSMs calculated by Proteome Discoverer. Bioinformatic analysis was performed in R. Gene Ontology Enrichment (Biological Process) using Bioconductor DOSE36 and cluster.Profiler37 packages and an online web-based gene analysis toolkit (Cellular Compartments and node graph)38. Heat maps were plotted using the ggplot2 package. Volcano plots, heat maps, and principal component analysis (PCA) were plotted using the ggplot2 package. Differential expression analysis was conducted with the Bioconductor DESeq39 package, using the spectral count-based quantification Peptide to Spectral Matching (PSM) (26) as input, and evaluation of the enriched signature genes was performed with the gene set enrichment analysis (GSEA) algorithm (27), employing 1000 permutation and default parameters.

## Results

### UV-C irradiation induce genomic damage and p53 activation in iPSC-MSC

In order to study how UV-C radiation modifies the biogenesis and cargo of sEVs, Mesenchymal Stem Cells derived from induced Pluripotent Stem Cells (iPSC-MSC) were irradiated with three UV-C light (254 nm) energy doses: 0.001 J/cm2, 0.01 J/cm2 and 0.1 J/cm2 (Figure 1A). To validate our experimental design, we assessed the induction of DNA Damage Response (DDR) in the UV-C irradiated iPSC-MSC. The three UV-C radiation intensities applied on iPSC-MSC caused the number of H2AX-*gamma* positive nuclei to increase with different velocities (Figure 1B and C). In addition, nuclear translocation of p53 was observed for the three irradiation intensities (Figure 1D), again showing slightly different kinetics for each intensity. Nuclear p53 signal peaked at 10 hrs after 0.001 J/cm2 irradiation, between 6 and 10 hrs after 0.01 J/cm2 irradiation and as soon as 3h after 0.1 J/cm2 irradiation. Since the ser-15 residue in the p53 N-terminal transactivation domain is a well-known target of the kinases ATM and ATR, key regulators in DDR, we sought to further validate our results assessing the nuclear localization of S-15 phosphorylated p53 at 6 hrs post UV-C exposure (figure 1E), the moment when it is predominantly nuclear. In addition, post-translational modifications of p53 are known to influence its stability so we analyzed p53 protein level up to 24hs post-radiation. Interestingly, total levels of p53 showed a significant increase 24 hrs after 0.001 J/cm2 and 0.01 J/cm2 irradiation compared to not irradiated iPSC-MSC, while 0.1 J/cm2 irradiation intensity did not cause such rise (figure 1F). Together, these results support previous reports that indicate a dose-dependent DNA damage response with UV-C irradiation (18). In addition, the results shown in figure 1 suggest that increasing doses of UV-C light may elicit different intra-cellular responses, leading to different fate decisions. Thus, we next sought to gain insight on how the sEVs biogenesis-related genes are affected by genomic stress.

**Figure 1:**
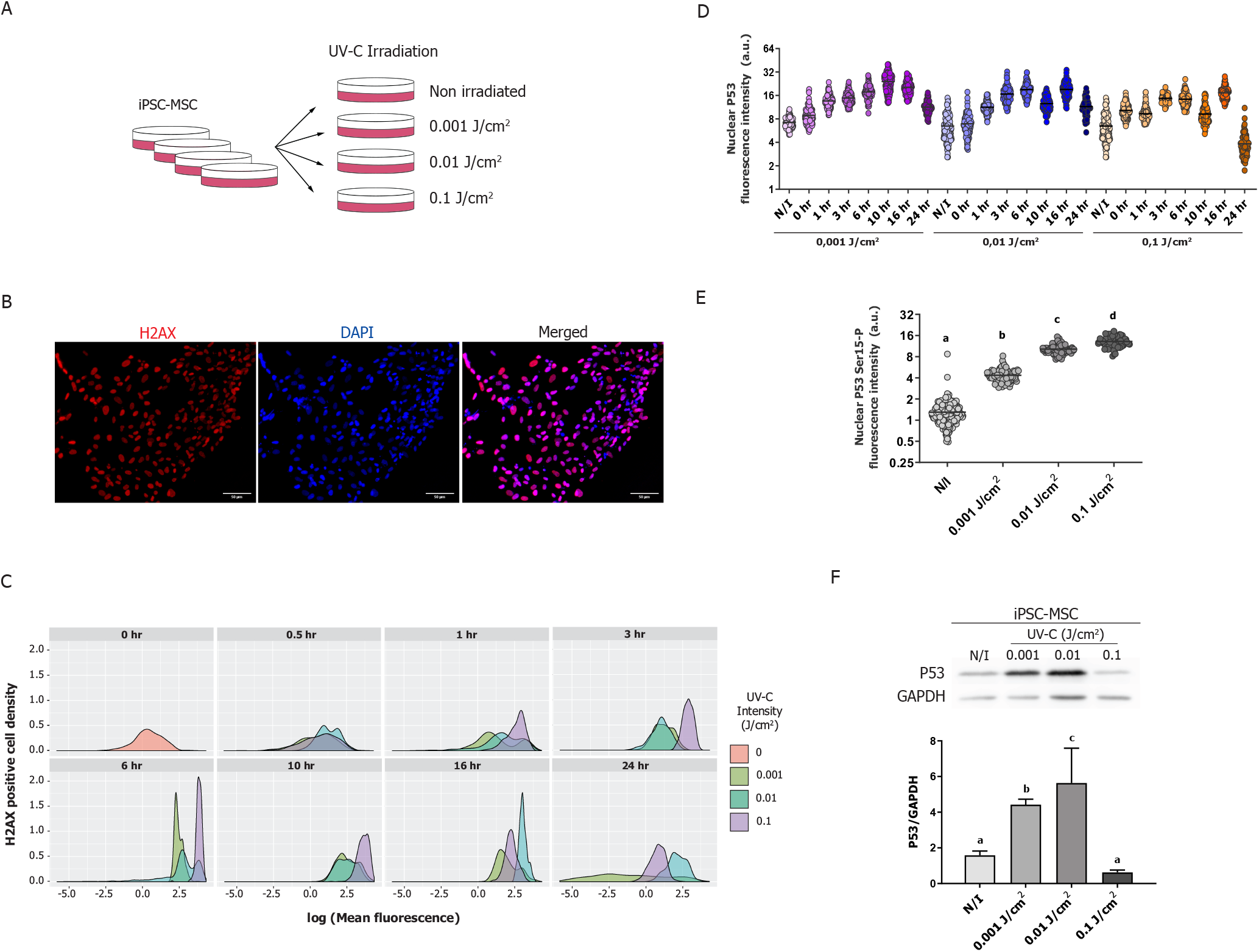
UV-C radiation causes genomic damage in iPSC-MSC. A) Experimental design used on iPS-MSC. Cells were irradiated or not (non irradiated) with three UVC light (254 nm) energy doses (0.001 J/cm2, 0.01 J/cm2 and 0.1 J/cm2) and both cells and supernatants were collected separately for downstream analysis. B) Representative fluorescence microscopy images of γ-H2AX signal after 6 hrs of UVC light irradiation in iPS-MSC. Nuclei were stained with DAPI. C) Distribution of quantified γ-H2AX signal intensity from iPS-MSC after different UVC irradiation times. D) Nuclear P53 fluorescence intensity at different times post irradiation in iPSC-MSC. E) Nuclear P53 Ser15-P fluorescence intensity at 6 hrs after irradiation in iPSC-MSC. F) P53 expression levels at 24 hrs after radiation assayed by Western Blot (WB). Left panel: Representative WB; Right panel: quantification of three independent assays. Result is shown +/− SD.

### UV-C irradiation does not modulate exosomal biogenesis pathway-related genes in MSC

Previous evidence indicates a role of p53 in the modulation of exosomal secretion. Thus, we analyzed the expression levels of genes involved in exosomal biogenesis pathways. Figure 2A shows the RNA levels of *STEAP3*, *SDCBP (SYNTENIN)*, *PDCD6IP (ALIX)* and *RAB27A*. From these, only *SYNTENIN* and *RAB27A* expression seemed to be transiently modulated with the mildest, 0.001 J/cm2 UV-C dose, although this transient modulation of RNA levels was neither reflected in their protein levels (figure 2B) nor replicated in experiments with Wharthon’s Jelly MSC (Supplementary figure 1). The absence of induction in *STEAP3* RNA levels is particularly interesting since it is proposed that Steap3 mediates the p53-dependent increase in exosome secretion. Although after UV-C exposure p53 was indeed activated and translocated to the nucleus, Steap3 levels did not increase regardless of the UV-C dose applied. These findings were further replicated in UV-C-irradiated Wharton’s Jelly MSC and in iPSC-MSC treated with another DNA damage inductor: the Topoiso-merase 1 inhibitor Camptothecin (supplementary figure 1B). Moreover, *in silico* analysis using MEME Suit motif occurrence search tool FIMO (http://meme-suite.org/) (28, 29), found no match for TP53 (JASPAR core vertebrate’s databa motif MA0106.3) on the 5000 bp upstream Steap3 transcription start site. FIMO did find, however, a TP53 motif in *ALIX* and *RAB27A* 5000 bp upstream regulatory sequence. The fact that neither RNA nor protein levels of Steap3 increased with the upregulation of p53, together with the apparent absence of p53 binding motifs on STEAP3 upstream sequence argues against a direct regulation of STEAP3 by p53 in our experimental model.

**Figure 2:**
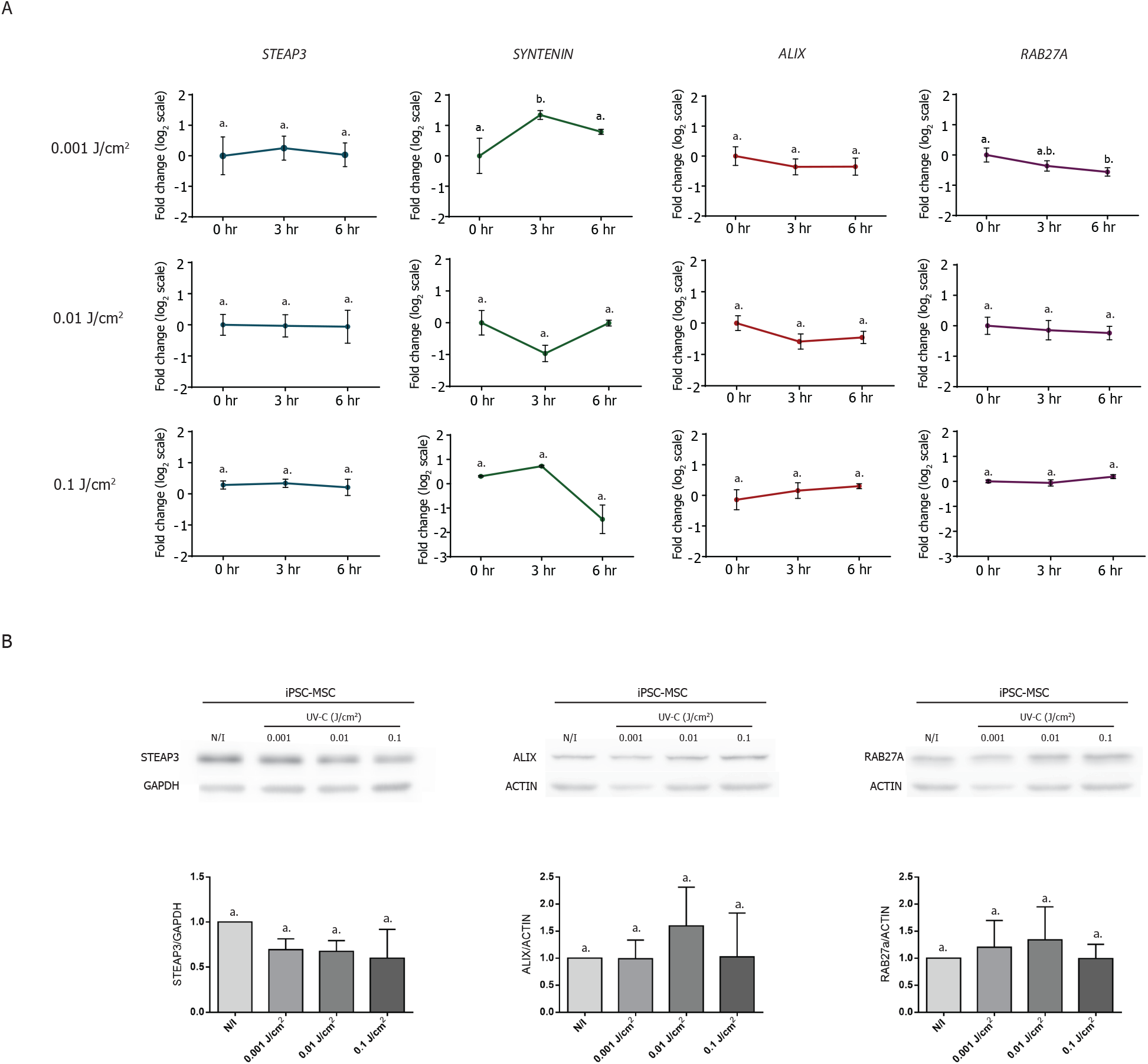
UV-C irradiation does not change the expression levels of genes and proteins involved in sEVs biogenesis pathway. A) RNA expression levels of *STEAP3*, *ALIX*, *SYNTENIN*, and *RAB27A* measured by qPCR in irradiated (3 and 6 hrs) and non irradiated (0 hrs) iPS-MSC. B) Analysis of protein expression levels of STEAP3, ALIX, and RAB27A assayed by Western Blot 8 hrs after irradiation in iPSC-MSC. Upper panel: Representative WB; Lower panel: quantification of three replicate assays.

### UV-C irradiation does not influence the amount of sEVs secreted

As we observed that expression of important genes involved in exosomal biogenesis did not change under genomic stress, we examined the number of sEVs secreted by irradiated and non irradiated iPSC-MSC. sEVs from 0.1 J/cm2 irradiated iPSC-MSC were not included in this analysis due to the high amount of cellular death evidenced when MSC treated with this dose were cultured with defined medium for conditioning purposes. Therefore, sEVs from non-irradiated, 0.001 J/cm2 and 0.01 J/cm2 were isolated from serum-free conditioned media using size exclusion chromatography. Isolated EVs were first assessed for the presence of CD63, CD81 and CD9 by flow cytometry (Figure 3A) and then profiled by Tunable Response Pulse Sensing (TRPS) (Figure 3B). In agreement with the results previously discussed, the number of secreted exosomes did not show substantial differences among UV-C doses. Interestingly, slight differences in size dispersion were observed in 0.01 J/cm2 exosomes compared to non-irradiated and 0.001 J/cm2.

**Figure 3:**
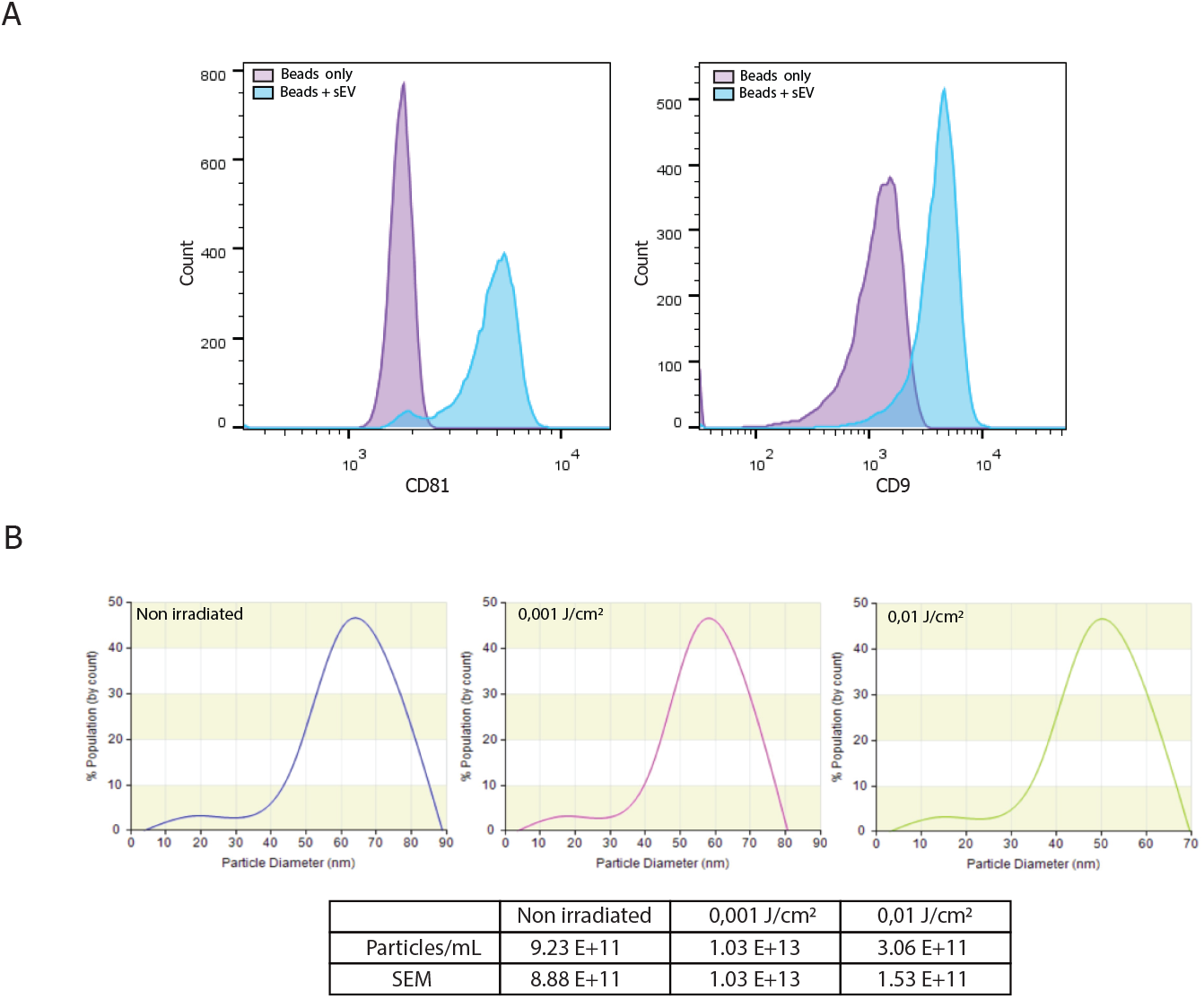
Isolation and quantification of sEVs. A) Flow cytometry of sEVs isolated using SEC columns. Vesicles were incubated with anti-CD63 magnetic beads followed by staining with CD9-APC-conjugated and CD81-PE-conjugated antibodies. B) Upper panel: quantification and size distribution of irradiated and non-irradiated iPSC-MSC derived sEVs assessed by Tunable Resistive Pulse Sensing (TRPS); Lower panel: nanoparticle count of three independent assays. Result is shown as mean +/− SEM.

### UV-C irradiation affects the functionality and composition of iPSC-MSC derived sEVs

As shown previously, sEVs secreted by iPSC-MSC are able to promote cell migration (10). Thus, we wondered whether this regenerative potential of MSCs-derived sEVs is maintained after cell irradiation. In order to evaluate this, we performed a wound healing Assay in which WJ-MSCs were exposed to sEVs obtained from irradiated and non-irradiated iPSC-MSCs. As expected, a significant improvement in wound closure was detected for sEVs from non-irradiated iPSC-MSCs relative to the control condition without sEVs (Figure 4A and B). Notably, this effect appears to be lost when sEVs from 0.001 J/cm2 and 0.01 J/cm2 UV-C irradiated cells were added to the culture medium. In particular, sEVs derived from 0.01 J/cm2 UV-C-treated MSC have no effect over wound closure at all. To assess if the cell cycle is altered in EVs-targeted cells, we stained them with propidium iodide after exposure to sEVs from non-irradiated, 0.001 J/cm2 and 0.01 J/cm2-irradiated cells. As seen in figure 4C, after 16 hours there is little to no variation in cell cycle phases. However, there seems to be a slight increase in s-phase duration for cells exposed to irradiated derived sEVs and a minor decrease in G1-phase duration for cells exposed to non-irradiated sEVs, though these variations were not statistically significant. Moreover, CyQuant assay revealed a decreased number of cells incubated with non-irradiated EVs after 14 hours. This decrease, in turn, diminishes when cells are exposed to irradiated sEVs (Figure 4D). Overall, these results suggest that MSCs sEVs pro-migratory properties are lost upon irradiation reinforced by the observation that wound closure alterations were not a byproduct of an increased cell number during the assay.

**Figure 4:**
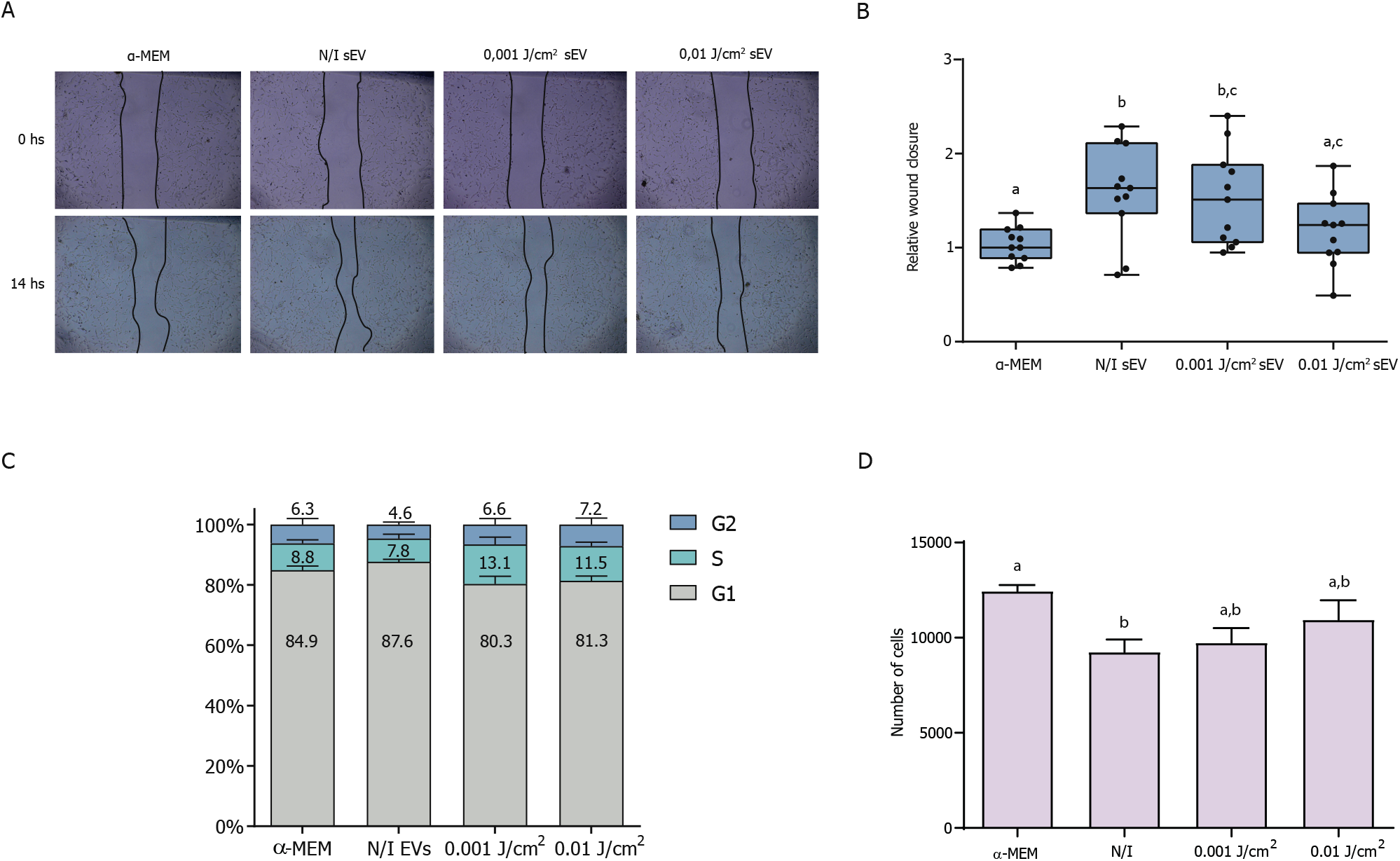
UV-C irradiation affects the pro-migratory properties of iPSC-MSC-derived sEVs. A) Representative wound healing assay. A WJ-MSC monolayer was scratched and incubated during 14 hrs with sEVs secreted by irradiated or non irradiated (N/I) iPSC-MSC, or without sEVEs (α-MEM). B) Quantification of cell-free areas as a measure of wound closure. Result is shown as mean +/− SEM. C) Cell cycle analysis by propidium iodide incorporation assayed by flow cytometry of iPSC-MSC exposed to irradiated or non-irradiated iPSC-MSC derived exosomes during 14 hrs. D) Cell proliferation analysis of WJ-MSC exposed to irradiated or non-irradiated iPSC-MSC derived exosomes during 14 hrs.

### UV-C irradiation affects composition of iPSC-MSC derived sEVs

Last, we sought to evaluate if the loss of function previously described could be explained due to a change in sEVs composition upon exposure of iPSC-MSC to UV-C light. In a previous work, we analyzed the proteomic content of iPSC, iPSC-MSC and WJ-MSC sEVs and found that, during differentiation towards MSC, iPSC sEVs acquire a specific stromal repertoire that explain iPSC-MSC sEVs’ regenerative properties. Therefore, we hypothesize that loss in migratory-induction ability of sEVs may be driven by a change in its proteomic cargo. Hence, we performed a proteomic analysis of sEVs from non-irradiated, 0.001 and 0.01 J/cm2 irradiated iPSC-MSC. As shown in figure 5A and supplementary table 2, we found that sEVs content changed substantially, with only 2 unique Uniprot IDs for sEVs derived from control iPSC-MSCs, 19 for sEVs from 0.001 J/cm2 radiated MSCs and 325 Uniprot IDs that exclusive to sEVs from 0.01 J/cm2 irradiated cells. In addition, we found that sEVs from 0.001 and 0.01 J/cm2 irradiated cells shared 144 Uniprot IDs while 37 Uniprot IDs were shared amongst the three conditions. Both Pearson’s Correlation Analysis and Principal Component Analysis showed a marked grouping amongst biological replicates of each condition but was able to distinguish between experimental conditions (Figure 5B and C).

**Figure 5.**
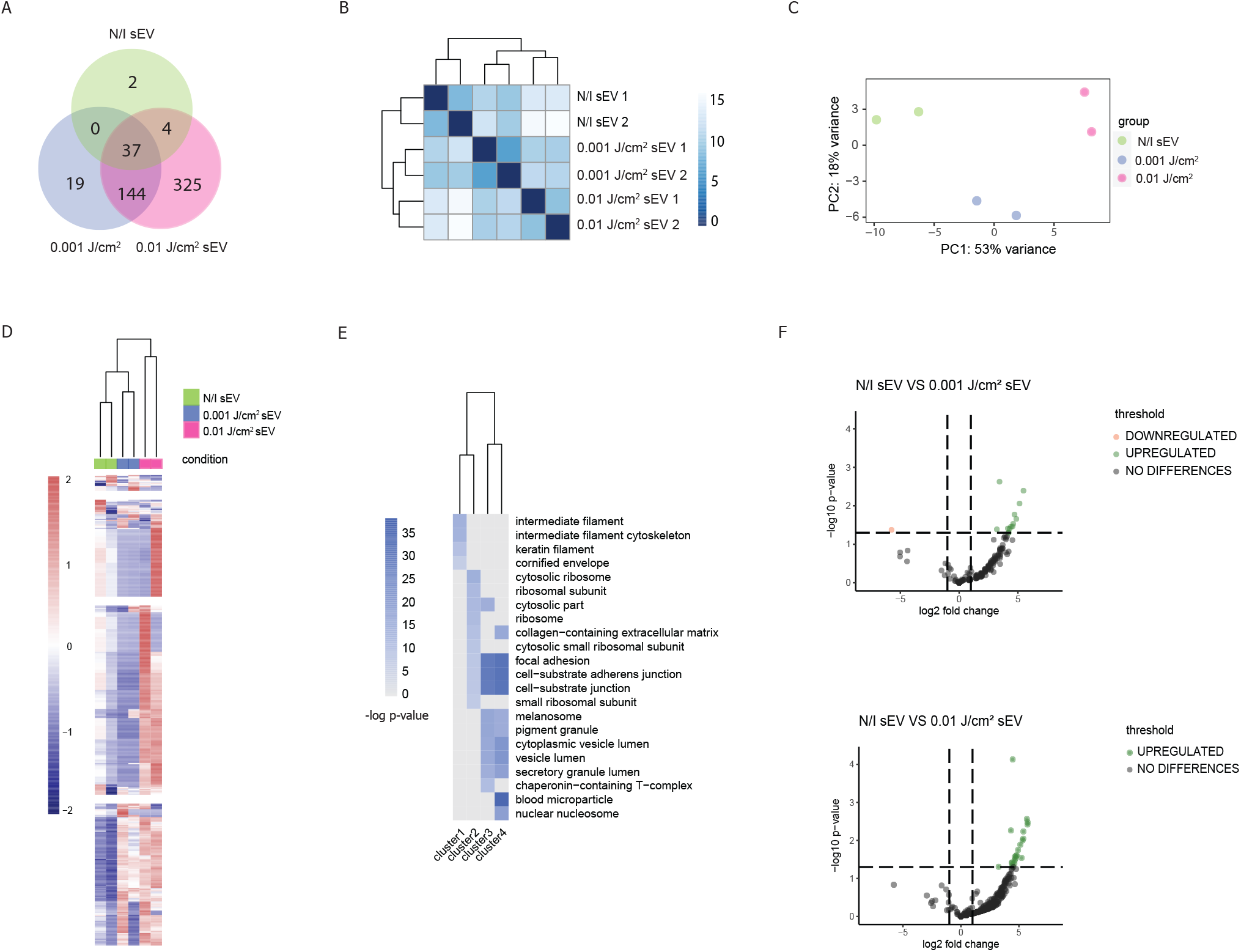
EVs’s content changes in response to genomic damage caused by UV-C radiation. A) Venn diagram of UniProt IDs identified by LC-MS/MS in non-irradiated, J/cm2 and 0.01 J/cm2 irradiated iPSC-MSC derived sEVs. B) Distance Matrix showing Parson’s coefficient between proteins in EVs derived from non-irradiated, 0.001 J/cm2 and 0.01 J/cm2 irradiated iPSC-MSC. Matrix was plotted using two replicates for each EV source. C) Principal component analysis showing the grouping of samples shown in A according to the variables that further explain their differences. D) Heatmap of protein abundance of non-irradiated, 0.001 J/cm2 and 0.01 J/cm2 irradiated iPSC-MSC derived sEVs. E) Gene Ontology analysis (Cellular Components) of clusters 1 to 4 UniProt IDs. F) Volcano plot representing the Differential Expression Analysis results of non-irradiated versus 0.001 J/cm2 and non-irradiated versus 0.01 J/cm2.

A Heatmap plot for protein abundance of non-irradiated, 0.001 J/cm2 and 0.01 J/cm2 irradiated iPSC-MSC derived sEVs revealed four distinct clusters (Figure 5D and Supplementary table 2). In particular, cluster 3 and 4 revealed a set of proteins that were over-represented only in 0.001 J/cm2 or both in 0.001 J/cm2 and 0.01 J/cm2 sEVs with respect to the non-irradiated condition, respectively. Gene ontology of these two clusters show that over-represented peptides belong to certain categories of cellular components such as focal adhesion structures and cell-substrate junctions (Figure 5E), hinting to a possible role of these proteins in the loss of pro-migratory properties of non-irradiated sEVs.

To further analyse this, we performed a differential expression (DESeq) analysis comparing non-irradiated to 0.001 J/cm2 and 0.01 J/cm2 irradiated MSC derived sEVs, respectively (Figure 5F). In both analyses a subset of proteins appeared consistently over-represented in sEVs from irradiated cells (Table 1). This includes representatives of Actin and Tubulin family (*ACTC1*, *TUBB5*, *TUBB4B*), Integrins *(ITGB3*, *ITGA2B*), Apolipoproteins *(APOE*, *APOB-100*), Heat Shock Proteins (*HSPA8*, *HSP90*) and, notably, the proteins Filamin A, Thrombospondin-1 (*TSP1*) and Talin-1. These last three are noteworthy because they negatively regulate cell motility and migration by either binding with Actin filaments (Filamin A), associating with integrins (Talin-1) or by being an integral part of the extracellular matrix (*TSP-1*). These proteins have been extensively characterized in cancer, however their role in stromal homeostasis and cell regeneration have been somewhat unexplored.

**Table 1:** UniprotID, Protein name and gene name of differentially expressed (DESeq) proteins between non-irradiated and 0.001 J/cm2 iPSC-MSC derived sEVs (NI vs 0.001 J cm2, top panel), and between non-irradiated and 0.01 J/cm2 iPSC-MSC derived sEVs (NI vs 0.01 J cm2, bottom panel).

| NI vs 0.001 J cm<sup>2</sup> |  |  |
| --- | --- | --- |
| UNIPROT ID | PROTEIN NAME | GENE NAME |
| P01023 | Alpha-2-macroglobulin | A2M CPAMD5 FWP007 |
| P02649 | Apolipoprotein E (Apo-E) | APOE |
| P04114 | Apolipoprotein B-100 | APOB |
| P05106 | Integrin beta-3 | ITGB3 GP3A |
| P07437 | Tubulin beta chain | TUBB TUBB5 OK/SW-cl.56 |
| P08133 | Annexin A6 | ANXA6 ANX6 |
| P08514 | Integrin alpha-IIb | ITGA2B GP2B ITGAB |
| P11142 | Heat shock cognate 71 kDa protein | HSPA8 HSC70 HSP73 HSPA10 |
| P14618 | Pyruvate kinase | PKM OIP3 PK2 PK3 PKM2 |
| P21333 | Filamin-A | FLNA FLN FLN1 |
| P68371 | Tubulin beta-4B | TUBB4B TUBB2C |
| P98160 | Basement membrane-specific heparan sulfate proteoglycan core protein | HSPG2 |
| Q9Y490 | Talin-1 | TLN1 KIAA1027 TLN |

| NI vs 0.01 J cm<sup>2</sup> |  |  |
| --- | --- | --- |
| UNIPROT ID | PROTEIN NAME | GENE NAME |
| P21333 | Filamin-A | FLNA FLN FLN1 |
| Q9Y490 | Talin-1 | TLN1 KIAA1027 TLN |
| P08133 | Annexin A6 | ANXA6 ANX6 |
| P08514 | Integrin alpha-IIb | ITGA2B GP2B ITGAB |
| P35579 | Myosin-9 | MYH9 |
| P01024 | Complement C3 | C3 CPAMD1 |
| P14618 | Pyruvate kinase | PKM OIP3 PK2 PK3 PKM2 |
| P07437 | Tubulin beta chain | TUBB TUBB5 OK/SW-cl.56 |
| P68032 | Actin, alpha cardiac muscle 1 | ACTC1 ACTC |
| P68371 | Tubulin beta-4B chain | TUBB4B TUBB2C |
| P07996 | Thrombospondin-1 (Glycoprotein G) | THBS1 TSP TSP1 |
| P02649 | Apolipoprotein E (Apo-E) | APOE |
| P01834 | Immunoglobulin kappa constant | IGKC |
| P04114 | Apolipoprotein B-100 | APOB |
| P04406 | Glyceraldehyde-3-phosphate dehydrogenase | GAPDH |
| P11142 | Heat shock cognate 71 kDa protein | HSPA8 HSC70 HSP73 HSPA10 |
| P08238 | Heat shock protein HSP 90-beta | HSP90AB1 HSP90B HSPC2 HSPCB |
| P05106 | Integrin beta-3 | ITGB3 GP3A |
| P01023 | Alpha-2-macroglobulin | A2M CPAMD5 FWP007 |
| P07355 | Annexin A2 | ANXA2 ANX2 ANX2L4 CAL1H LPC2D |

Altogether these results suggest that MSC irradiated sEVs carry a distinct and particular cargo, including proteins that may negatively regulate cell migration and that can explain the reverting of the pro-migratory capabilities of non-irradiated MSC sEVs.

## Discussion

DNA damage response triggers an activation in p53 that in turn elicits a myriad of biological effects that help the cells to cope with that damage. In particular, DSB is proposed to increase the secretion of EVs in a p53 dependent manner (19, 21–24). It has been proposed that this increased secretion requires the metalloreductase *STEAP3*, although the exact molecular mechanism is not yet well understood (20, 30, 31). Additionally, EVs from gamma-radiated and UV exposed cells are able to induce biologic effects on cells that were not exposed through a mechanism known as Radiation-induced bystander effects (RIBE) (32–34). Most of the evidence for these phenomena come from experiments in cancer cell lines and treated with ionizing radiation. However, little is known of the effect of UV-C light on mesenchymal stem cells and their extracellular vesicles.

In this work, we explored how UV-C radiation affects the biogenesis and constitution of sEVs in iPSC-MSC. We observed that as little as 0.001 J/cm2 of UV-C light exposure was enough to recruit H2AX to DNA, and induce and activate p53. However, different doses of UV-C light appear to elicit different molecular responses to DDR. Not only H2AX is activated with different kinetics (figure 1C), but p53 also seems to be translocated to the nucleus and stabilized differently (figure 1D). Moreover, p53 protein levels appear to diminish strongly 24 hrs after exposure to an extremely high 0.1 J/cm2 UV-C dose. These findings are in accordance with previous reports that UV-radiation induces dose-dependent regulation of p53 response (Latonen 2001). Noteworthy, MSC have a high resistance to UV light (Lopez Perez 2019), but the 0.1 J/cm2 UV-C dose resulted in cell death 24 hrs post-irradiation unless they are cultured with 10% platelet lysate.

One aspect of DDR we were particularly interested in was the consequence of genomic stress over MSC regenerative secretome. There is abundant evidence that stress increases the secretion of EVs. Moreover, past evidence supports that ionizing radiation augments exosome secretion caused by the upregulation of p53 (21**?**). Thus, we hypothesized that UV-C radiation dependent p53 upregulation would increase the levels of exosomal biogenesis related genes such as *ALIX*, *SYNTENIN* and *RAB27A*. Surprisingly, this was not the case. None of these genes showed upregulation of their mRNA levels up to 6 hrs post radiation or evidenced changes in their protein levels 8 hrs post radiation. In addition, *STEAP3*, a metalloreductase proposed to be a key link in the senescence p53 dependent secretory pathway (20), did not show transcriptional or post-transcriptional modulation neither at these same times (figure 2) nor at 24 hrs post radiation (data not shown). A more careful dissection of the *STEAP3* regulatory region by the bioinformatic tool FIMO predicted no p53 binding sequence in the 5 Kbp upstream region of this gene.

In addition, queries for TP53 target genes in ChIP-Atlas database (https://chip-atlas.org/) show some low-score p53 binding in a cell-line dependent manner and when the predicted *STEAP3* regulatory region is extended up to 10 Kbp. Taken together, this evidence suggests that p53, though active, is unable to activate *STEAP3* in our experimental model. Finally, the most compelling piece of evidence against a DDR modulation of sEVs secretion in MSC, is the direct measurement of nanoparticles in the MSC conditioned media. In our experience, particle count by TRPS (figure 3) of sEVs from irradiated and non-irradiated cells has been fairly stable, albeit with a slight increase in EVs secreted by 0.001 J/cm2-radiated cells.

Radiation-induced bystander effects (RIBE) is a well-established phenomenon accounting for biological effects exerted onto non-irradiated cells. Moreover, sEVs are known to convey multiple stress related biological effects over the cell niche (35, 36). In this work we confirmed that sEVs from iPSC-MSC are able to increase wound closure *in vitro* and we report a loss of that migration-inducing capacity upon cell radiation. In a previous work, we related iPSC-MSC’s sEV composition, rich in extracellular matrix peptides and growth factors, to their regenerative capabilities (10). Here, we describe that 0.001 and 0.01 J/cm2 UV-C radiated derived sEVs have a different protein composition than sEVs from non-irradiated cells, bearing an increased amount of cytoskeleton proteins and migration-inhibiting molecules. In particular, proteins that participate in focal adhesion and cell-substrate junctions like Filamin A, Talin-1 and Thrombospondin-1 seem to be upregulated in sEVs upon UV-C irradiation of cells. Interestingly, these three proteins are frequently related with tumor progression and tumoral cell migration leading to metastasis (37–39), but have been poorly studied in the physiological processes that promote tissue regeneration. Aging and other pathologies are known to alter MSC and their secretome (40), so studying how these changes are linked to a decrease in migration would be highly valuable for the design of new regenerative therapies. Last, the intimate association of Filamin A, Talin-1 and Thrombospondin-1 with oncology prompts the interesting question on whether MSC-EVs with a fine-tuned composition of proteins might be suitable for anti-metastasis therapies.

Altogether this work proposes that UV-C induced DDR causes different cellular effects in a dose-dependent manner and that those effects translate into a modulation of MSC derived sEVs cargo. This content modulation may produce, in turn, a change in sEVs effect over target cells in their niche, decreasing cell migration and hence diminishing their regenerative capacity.

## Acknowledgements

This work was supported by grants from the National Agency for Scientific and Technical Promotion (ANPCyT) and from the Scientific and Technical Research Fund (FONCyT) PICT-2015-0868. Authors would like to thank FLENI-CONICET and Pérez Companc Foundation for their continuous support.

## Supplementary Material

**Table S1.**
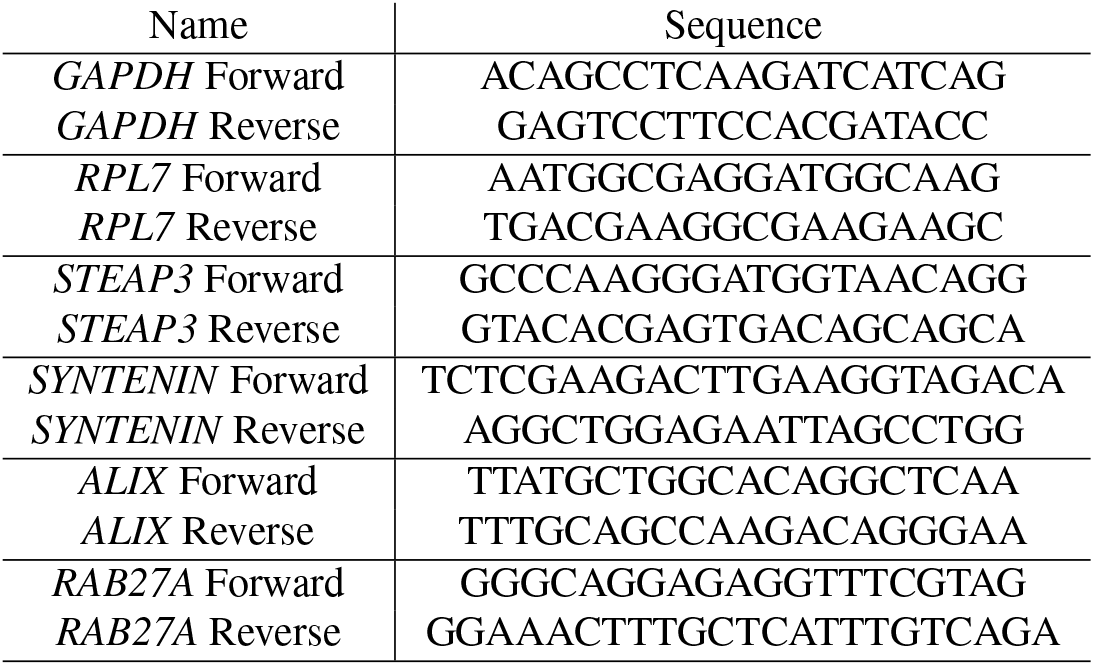
Primers sequences

**Supplementary Figure 1.**
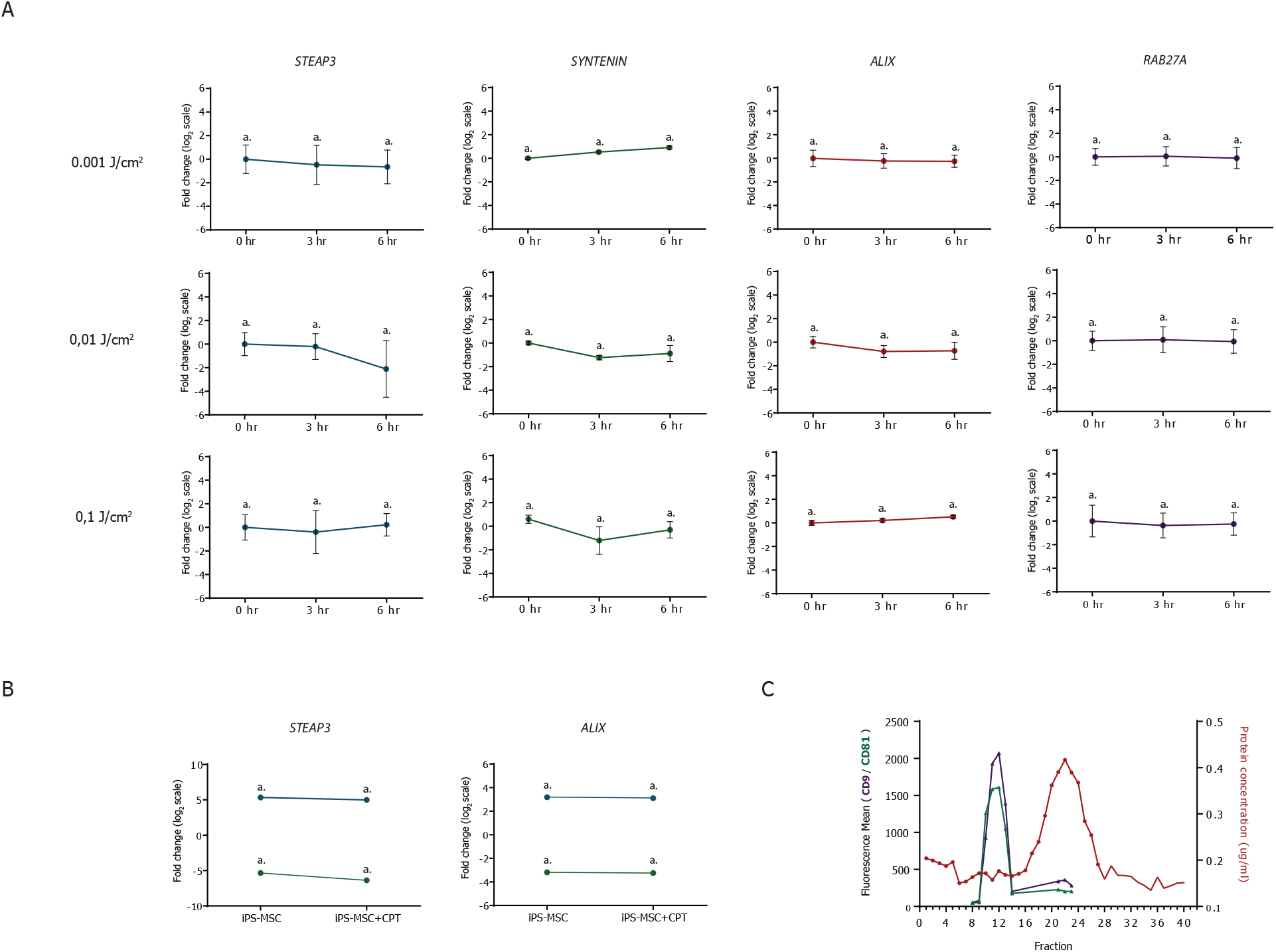
A) Analysis of mRNA expression levels of STEAP3, ALIX, SYNTENIN, and RAB27A quantified by RTq-PCR in irradiated and non irradiated WJ-MSC. B) Analysis of mRNA expression levels of STEAP3 and ALIX quantified by RTq-PCR in iPSC-MSC treated with the DNA damage inductor Topoisomerase 1 inhibitor Camptothecin. Each line represents a biological replicate. C) Protein concentration and fluorescence mean of CD9 and CD81 of 40 eluted fractions of the Size Exclusion Chromatography column. It can be distinguished how proteins present in conditioned medium are eluted separately from sEVs.

**Supplementary Figure 2.**
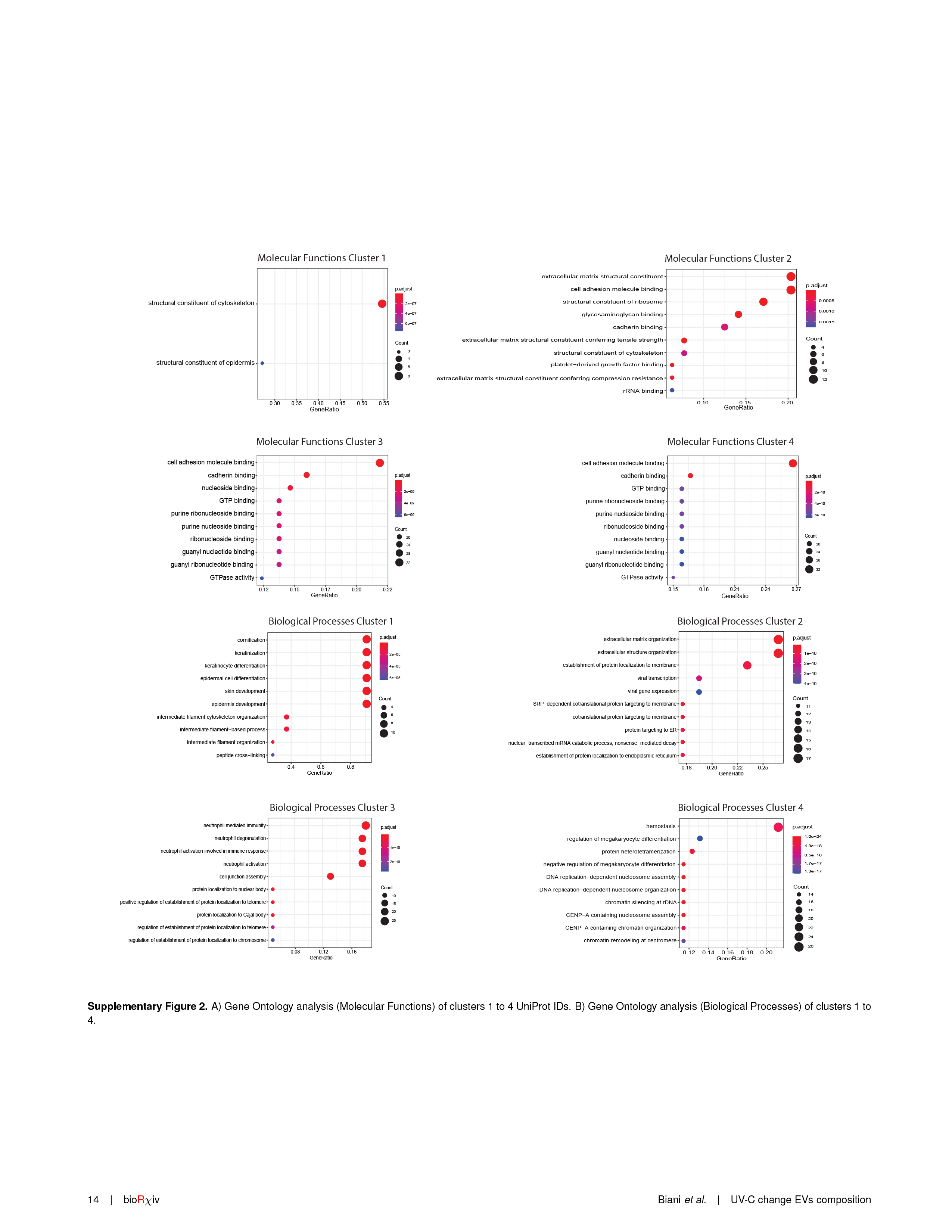
A) Gene Ontology analysis (Molecular Functions) of clusters 1 to 4 UniProt IDs. B) Gene Ontology analysis (Biological Processes) of clusters 1 to 4.

